# ExTaxsI: an exploration tool of biodiversity molecular data

**DOI:** 10.1101/2020.11.05.369983

**Authors:** Giulia Agostinetto, Anna Sandionigi, Adam Chahed, Alberto Brusati, Elena Parladori, Bachir Balech, Antonia Bruno, Dario Pescini, Maurizio Casiraghi

## Abstract

**Background:** The increasing availability of multi omics data is leading to continually revise estimates of existing biodiversity data. In particular, the molecular data enable to characterize novel species yet unknown and to increase the information linked to those already observed with new genomic data. For this reason, the management and visualization of existing molecular data, and their related metadata, through the implementation of easy to use IT tools have become a key point for the development of future research. The more users are able to access biodiversity related information, the greater the ability of the scientific community to expand the knowledge in this area.

**Results:** In our research we have focused on the development of ExTaxsI (Exploring Taxonomies Information), an IT tool able to retrieve biodiversity data stored in NCBI databases and provide a simple and explorable visualization. Through the three case studies presented here, we have shown how an efficient organization of the data already present can lead to obtaining new information that is fundamental as a starting point for new research. Our approach was also able to highlight the limits in the distribution data availability, a key factor to consider in the experimental design phase of broad spectrum studies, such as metagenomics.

**Conclusions:** ExTaxI can easily produce explorable visualization of molecular data and its metadata, with the aim to help researchers to improve experimental designs and highlight the main gaps in the coverage of available data.

## 1 Introduction

In recent years, studies investigating biodiversity at large scale started to create and incorporate molecular data. In particular, the spread of metagenomic studies (e.g. metabarcoding) have contributed to an exponential increase in genomic data availability. Thanks to this large amount of new information it is possible to expand our knowledge and enhance our scientific investigation capacity in many fields of research [47], ranging from macro-ecology and ecosystem monitoring, to food safety control, forensics applications and microbiome identification [15, 47, 52]. Different groups of researchers emphasized the wealth of information collected in biological and molecular databases, with the aim to improve usefulness and reusability of data [21, 39, 57]. Therefore, building experimental designs that consider the totality of the data present in the biological databases could certainly increase the efficiency of these studies, and lead to more robust results [1, 41].

Biodiversity data retrieval and exploration has become a big data issue, forcing researchers to use Information Technologies (IT) tools to manage those data. In particular, the interpretation of results derived from metagenomic approaches requires computational pipelines and IT infrastructures that improve over time, but are strongly linked to the availability of pre-existing data stored in online databases (e.g. ENA - www.ebi.ac.uk/ena; and NCBI - https://www.ncbi.nlm.nih.gov/).

Visualization remains an effective strategy not only to aggregate and present the research results, but also to guide advanced investigations [22, 29]. At this moment, reference databases where molecular and taxonomic data are friendly explorable and punctually updated exist only for few molecular markers, such as SILVA for 16S and 18S genes [48], BOLD for animals and plants [50] or UNITE for Fungi domain [44]. However, these data resources are not representative of all the genomic and taxonomic diversity collected to date. On the other hand, although GenBank still resumes the majority of genetic data and their related metadata currently available [3, 5, 30], such information is not always easy to access without specific bioinformatics skills, which is a limiting factor to a large audience of scientists.

With the aim to help biologists to improve experimental designs and to encourage data exploration and exploitation, we have developed a tool, ExTaxsI (Exploring Taxonomies Information), to facilitate the molecular data integration with its associated metadata, eventually retrieved from heterogeneous sources. Moreover, its ease of use interface will help researchers and practitioners in the visualization phase. ExTaxsI can both query NCBI Nucleotide database for molecular data and accept data from an external source, exploiting the standard taxonomy notation. The tool is linked to NCBI taxonomy database [17] and ETE toolkit [26], in order to produce standard formats readable by most common software that deal with taxonomic information [4, 7, 8, 38, 51, 56], such as QIIME2 platform [7]. The tool is applicable to any marker, gene name or taxonomic group, so it is possible to create non-standard marker genes database usable in metagenomic/metabarcoding taxonomic assignment tools [7]. In addition, thanks to the integration of the NCBI query tool [11], ExTaxsI can reorganize personal datasets in a standardized format in order to easily describe taxonomic variability and geographic provenance of records.

## 2 ExTaxsI

ExTaxsI is a bioinformatic tool aimed to elaborate and visualize molecular and taxonomic information via a simple interface. This open-source user friendly instrument, developed in Python 3.7, starting from a list of taxa or gene name/s (as illustrated in Figure 1), allows i) the search of taxonomic, genetic and biogeographical data through NCBI databases, ii) the creation of local and formatted nucleotide sequences (FASTA format) dataset and iii) their related taxonomy classification paths/datasets, thanks to the integration of NCBI taxonomy data, iv) the creation of genetic markers lists coming from different studies and finally v) the production of interactive plots starting from NCBI query search results or directly from offline taxonomic files, including representative graphs for the exploration of taxonomy and refinement of biogeographical data, creating geographical maps with the locations of the species analyzed (Figure 1). It is important to note that ExTaxsI outputs are compatible with other tools for taxonomic assignment purposes such as QIIME2 platform [7].

**Figure 1.**
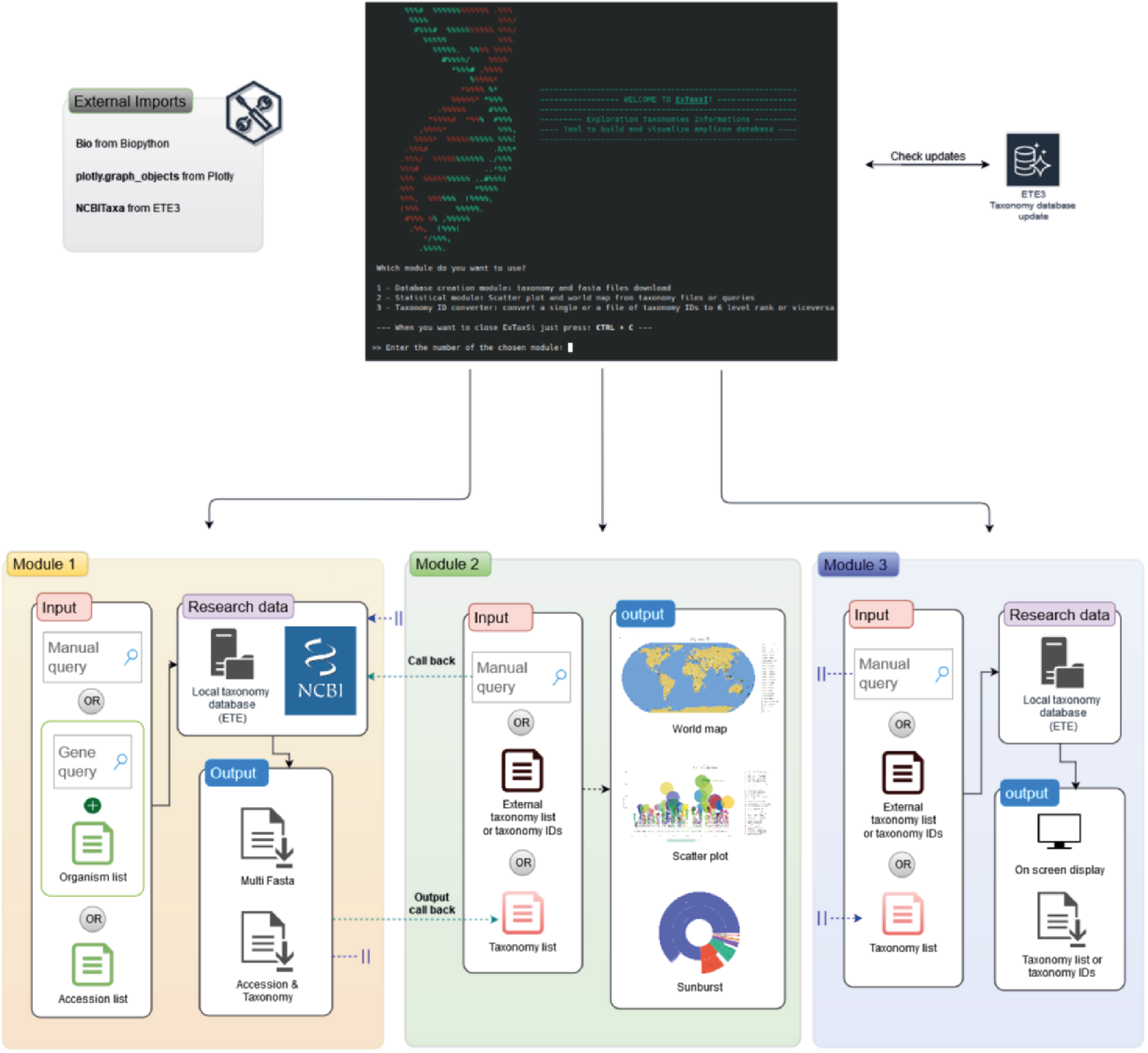
ExTaxsI pipeline: module 1 (orange) searches and creates files and databases; module 2 (green) processes georeferenced or taxonomic data for the creation of graphs and plots; module 3 (blue) converts taxonomic data into taxonomic ID (TaxID) and vice versa.

The communication with NCBI server is mediated by the Entrez module [11], implemented in Biopython library [10], which allows to search, download and parse query results. To help NCBI interaction, when the requests are less than 2500, the search key is composed by a single query, otherwise the query will be splitted in groups of 2500 generating temporary files, which are then merged into single output file at the end of the process.

Regarding taxonomy handling, ETE toolkit was exploited [26]. In particular, ETE allows to create and maintain a local taxonomy database up to date by extrapolating the 6 main ranks (phylum, class, order, family, genus and species). If the organism is poorly described or it is an unknown species, the Taxonomy ID (i.e. TaxID) of its ancestor (known as parent TaxID) in ETE taxonomic tree is then used and converted into its scientific correspondent name. It is important to underline that all queries are carried out locally, avoiding unnecessary delays and allowing the integration of the tool in genomic and metagenomic pipelines.

Finally, the extracted data are visualized through the ScatterPlot and the SunBurst (Expansion Pie) for the taxonomy exploration, and ScatterGeo for the geographic metadata plotting.

### 2.1 Use cases

Being ExTaxsI a taxonomy focused data exploration tool, we designed three possible scenarios of increasing complexity, to challenge it with increasing taxonomic variability and dimension of accession entries. The first scenario hypothesizes a query to explore data with i) low taxonomic variability and a high number of expected entries (1 species, more than 300,000 entries). The second scenario provides ii) a high taxonomic variability and a large expected number of entries (about 500 species, more than 300,000 entries). The third and more complex scenario explores a iii) complete case study with taxonomic input intersected by molecular data. As case studies of the first two scenarios, we focused on taxa of interest in marine fisheries: 1) the cod fish species (*Gadus morhua*), for which a worldwide economic interest exists, and 2) its taxonomic group at order level - the *Gadiformes* order - which supports long-standing commercial fisheries and aquaculture. These two case studies evaluate the capacity to explore data and to fill the geographic distribution of a species, prospecting also the available genes information to perform a genetic survey (e.g. DNA metabarcoding study). With the third use case, we aimed at demonstrating the flexibility of ExTaxsI in different contexts: a genetic exploration of the available data in NCBI associated to *SARS-CoV-2* virus - a very recent topic that involved many research groups, leading to huge amounts of data collected and deposited in public sources [6]. A large scale exploration of data related to this topic can potentially improve the reliability of results and can provide valuable evidence to inform decisions on public health protection, both now and most importantly in the future.

#### 2.1.1 Insights into two taxonomic groups of commercial interest

The first scenario is the case of *Gadus morhua* species, also called Atlantic cod. In detail, *Gadus morhua* is a large, cold-adapted teleost fish that supports long-standing commercial fisheries and aquaculture [27, 28, 33, 34, 54].

ExTaxsI retrieved a total of 366,963 accessions using the taxonomy ID through the following query: “txid8049[ORGN]” (where 8049 is the specific *Gadus morhua* TaxID; 18 of June, 2020). Only 53,695 entries showed a ‘gene’ tag investigable by ExTaxsI. As a unique species, we decided to represent the results obtained from genes survey (Figure 2) and the world map plot (Figure 3). Regarding gene distribution, the most abundant gene is CYTB (with 985 accessions), followed by COI (434) and ND2 (311). The remaining most abundant genes are the other ND portions and Cytochrome Oxidase fragments (COIII and COII), belonging to the mitochondrial genome. These results show the increased effort in sequencing “standard” barcoding markers, while moderately sequencing whole mitochondrial genomes. The remaining genes in the retrieved list and their relative accession frequencies distribution (see the complete list in Additional file 1) demonstrate that the entire genome of this species was sequenced). These results are in line with those obtained by Knudsen and colleagues (2019), where they personally developed specific primers for CYTB amplification, as it is a widely used marker in fish molecular characterization.

**Figure 2.**
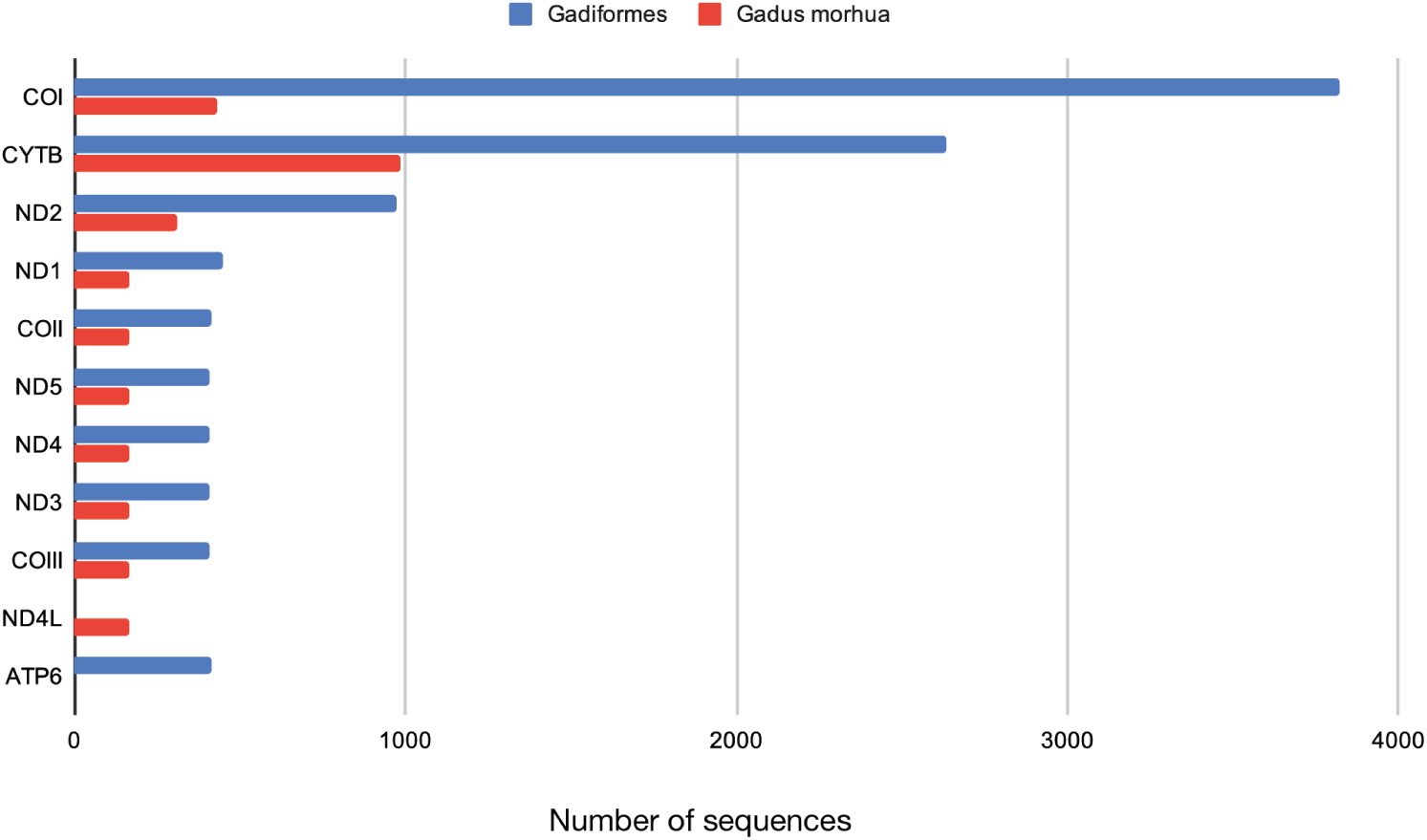
Gene distribution of accessions with complete ‘gene’ tag among *Gadus morhua* and *Gadiformes* taxons.

**Figure 3.**
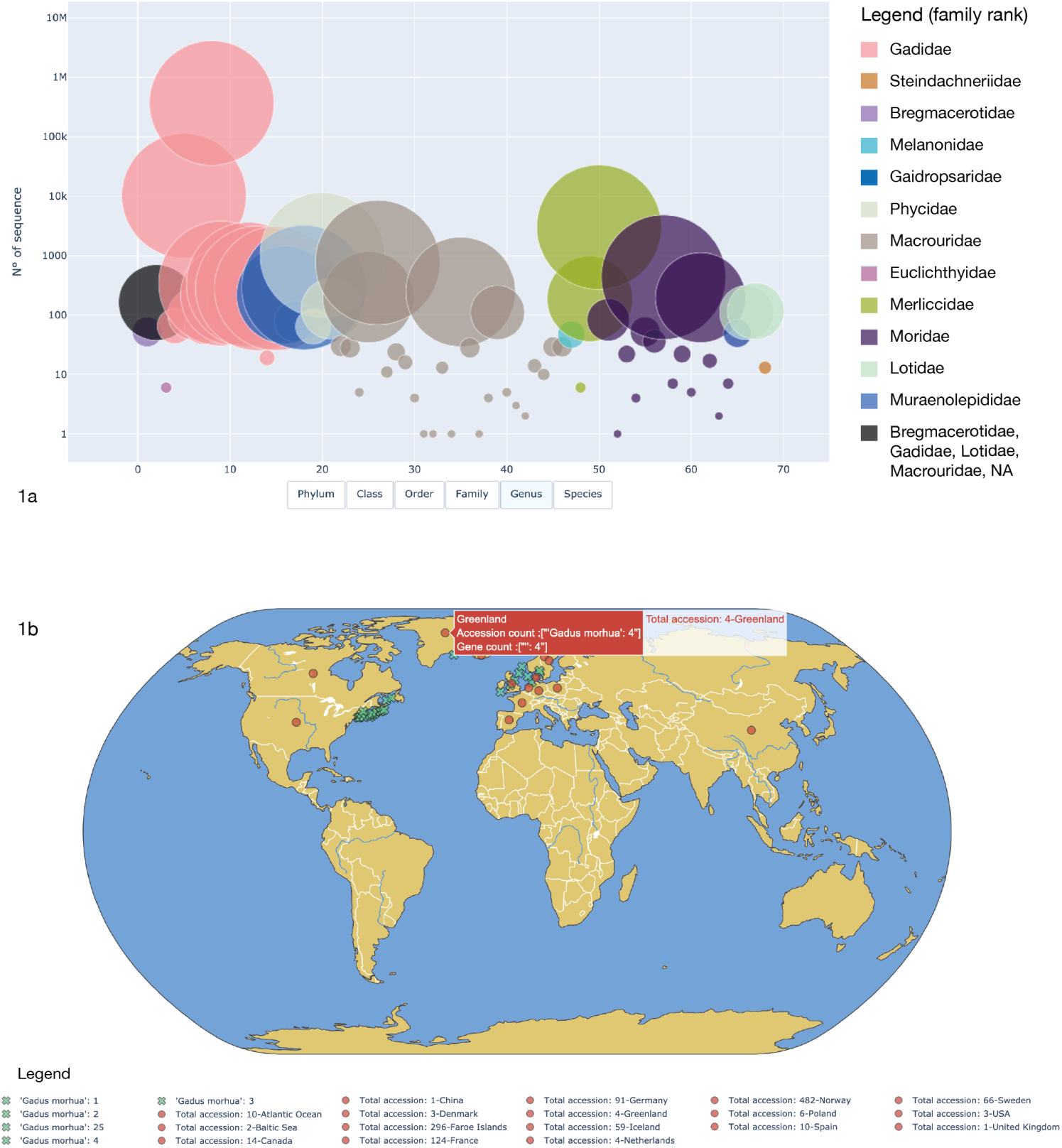
3a) Scatter plot of *Gadiformes* accessions to represent sequence abundances among families; 3b) World map plot of *Gadus morhua* distribution considering geographic metadata extracted from records.

Regarding the geographic area, the *Gadidae* family has a circumpolar distribution, comprising species occurring principally in northern and cool seas [28]. Further, as reported by Jorde and colleagues (2018), in Norway we can recognize four distinct stocks of the Atlantic cod: (1) the oceanic Northeast Arctic cod, (2) coastal cod north of 62°N, (3) coastal cod south of 62°N, and (4) a North Sea/Skagerrak stock, the most densely populated region in Norway [28]. This geographic distribution is partly visible via the metadata extracted by ExTaxsI, as shown in the world map plot in Figure 3b (Additional file 2).

The second scenario takes as an example the *Gadiformes* Order (phylum: *Chordata*; class: *Actinopterygii*), a major group of organisms belonging to marine fisheries. It includes many important food fishes, variously marketed as cods, hakes, grenadiers, moras, moray cods, pelagic cods, codlets and eucla cods [43]. As a vast group, it comprises more than 500 species, which contribute to more than a quarter of the world’s marine fish catch [13, 43].

Via ExTaxsI, this Order was explored using the following query “txid8043[ORGN]”, yielding 388,603 accessions (where 8043 is the specific *Gadiformes* TaxID; 22 of June, 2020), where 60,703 showed the ‘gene’ tag. As a group spread on different taxonomic levels, both taxonomy and gene reports were created. In detail, in order to explore taxa distribution and accessions abundances across the entire Order, the tool created ScatterPlot and SunBurst HTML outputs. In Figure 3a genus abundances are documented in ScatterPlot modality, while SunBurst and entirely interactive plots are available in the Supplementary Material section (Additional files 3 and 4).

As it is shown, *Gadidae* is the most abundant family, considering the number of accessions available. In fact, a total of 380,658 accessions populate this group, followed by *Merlucciidae* (3,196) and *Macrouridae* (1,581) families. These results are in accordance with the literature, a *Gadidae* family is a primary marine, bottom-dwelling family of fishes in the Order of *Gadiformes* with great commercial power [33, 43].

Further, considering the ScatterPlot in Additional file 3, the interactive visualization allowed us to explore the taxonomy distribution among the accessions available, changing dynamically the rank that we want to explore. This feature allows us to disclose that the genus *Gadus* is the most abundant of the entire dataset, highlighting that *Gadus morhua* species composed 94,43% of all the data. This is an expected result, as *Gadus morhua* is documented to be a key species both in the North Atlantic ecosystem and commercial fisheries, with an increasing aquaculture production in several countries [28]. Considering the genetic information reached by ExTaxsI, a total of 28,839 unique genes were found from the 60,703 completely tagged accessions. A classification of the most ten abundant genes is reported in Figure 2. As shown in the figure, at the first position we can find the COI gene, a widely used marker gene in metabarcoding projects (Knudsen et al. 2019), that deal mainly with animals detection [47], followed by CYTB and ND2 [47].

Concluding with these two case studies, the tool was able to accurately portrait the state of the art of the genetic information available in NCBI. Comparing the most abundant genes found among the records, it is possible to see a thin discrepancy between the two taxa explored (Figure 2), highlighting the disclosures that the survey can report. In general, the detection of mitochondrial genes, coding for Cytochrome Oxidase subunit I (COI) and Cytochrome B (CYTB), is in accordance with the reliability of these DNA barcodes, principally used in the discrimination of animal species [23, 24, 42]. To date, considering the subjects of our case studies, diverse studies have used COI or CYTB barcoding to identify seafood products and explore broad patterns in fish mislabelling [9, 16, 18, 40, 49, 59, 61].

Regarding the extraction of geographic metadata from NCBI records, the completeness and collection of data can improve drastically the biogeographic and ecological research, allowing not only to explore sampling areas, but also to improve phylogeography investigations, biodiversity monitoring and environmental genomics strategies [12, 47].

The unbalance between the number of records and the number of genes explorable is in some cases due to the incompleteness of the ‘gene’ tag. In the very recent years genome sequences started playing a key role into public repositories, making sequences available for sharing and reuse. Submission process can be challenging and errors can affect the availability of the data. For this reason, there is a wide interest to integrate standardized procedures into the annotation process [19]. The promotion of FAIR principles and best practices can certainly avoid the error propagation in sequence databases [46, 58], making the data fully explorable in the future.

#### 2.1.2 Explore biodiversity data in pandemic outbreak: the case of SARS-CoV-2

The severe acute respiratory syndrome coronavirus 2 (*SARS-CoV-2*) is an enveloped, positive-sense, single-stranded RNA virus that causes coronavirus disease 2019 (COVID-19). RNA and structural proteins are included into virus particles and mediate host cells invasion. After cell infection, RNA encodes structural proteins that make up virus particles. Virus assembly, transcription, replication and host control are mediated by nonstructural proteins [36]. The pandemic linked to *SARS-CoV-2* highlighted hidden virus reservoirs in wild animals and their potential to occasionally spillover into human populations [36]. A detailed understanding of this process is crucial to prevent future spillover events. As reported in the seminal paper of Andersen and colleagues (2020) [2], the risk of future re-emergence events increases if *SARS-CoV-2* pre-adapted in another animal species. *SARS-CoV-2* probably originated from *Rhinolophus affinis* bats, with pangolin (*Manis javanica*) as intermediate host [2]. Recently, other animal species were supposed to be possible intermediate hosts in between bats and humans. To date, ACE2 (Angiotensin-converting enzyme 2), the receptor which binds to the receptor-binding domain (RBD) of *SARS-CoV-2* S protein [35], is reported as crucial in host invasion.

To test our approach and explore the genetic information available in NCBI, we decided to extrapolate information of the ACE2 gene from the *Vertebrata* taxonomic group, with the following query: “txid7742[ORGN] AND ACE2[gene]” (where 7742 is the specific *Vertebrata* TaxID; 28 of June, 2020). The results show that the ACE2 gene is widely distributed throughout *Vertebrata*: we obtained a total of 1,189 accessions, distributed mainly among the *Mammalia* Class, with a high representation in *Actinopteri* and *Aves* groups (Figure 4; Additional files 5 and 6 are provided for an interactive exploration). In details, *Primates, Rodentia* and *Chiroptera* are the most represented, with 239, 132 and 108 accessions respectively. Supporting the exploitation of molecular data survey, Luan and colleague (2020) [37] analyzed the affinity to S protein of the 20 key residues in ACE2 from mammal, bird, turtle, and snake, and suggested that *Bovidae* and *Cricetidae* should be included in the screening of intermediate hosts for SARS-CoV-2. In addition, thanks to the analysis of spike glycoprotein sequences from different animals, the study of Dabravolski and Kavalionak [14] suggested that the human *SARS-CoV-2* could also come from yak as an intermediate host. ExTaxsI has the advantage to provide the complete list of taxa, allowing an exhaustive exploratory research. It allows to download all the sequences available for the query input, generating in turn the input for downstream analyses, such as the calculating of sequence similarities among different taxa. Further, investigating shared features with other species can have important implications for understanding potential natural reservoirs, zoonotic transmission, and human-to-animal transmission. Noteworthy, the survey can give researchers an instrument to download data with a user-friendly approach, exploring interactively the data and program experiments.

**Figure 4.**
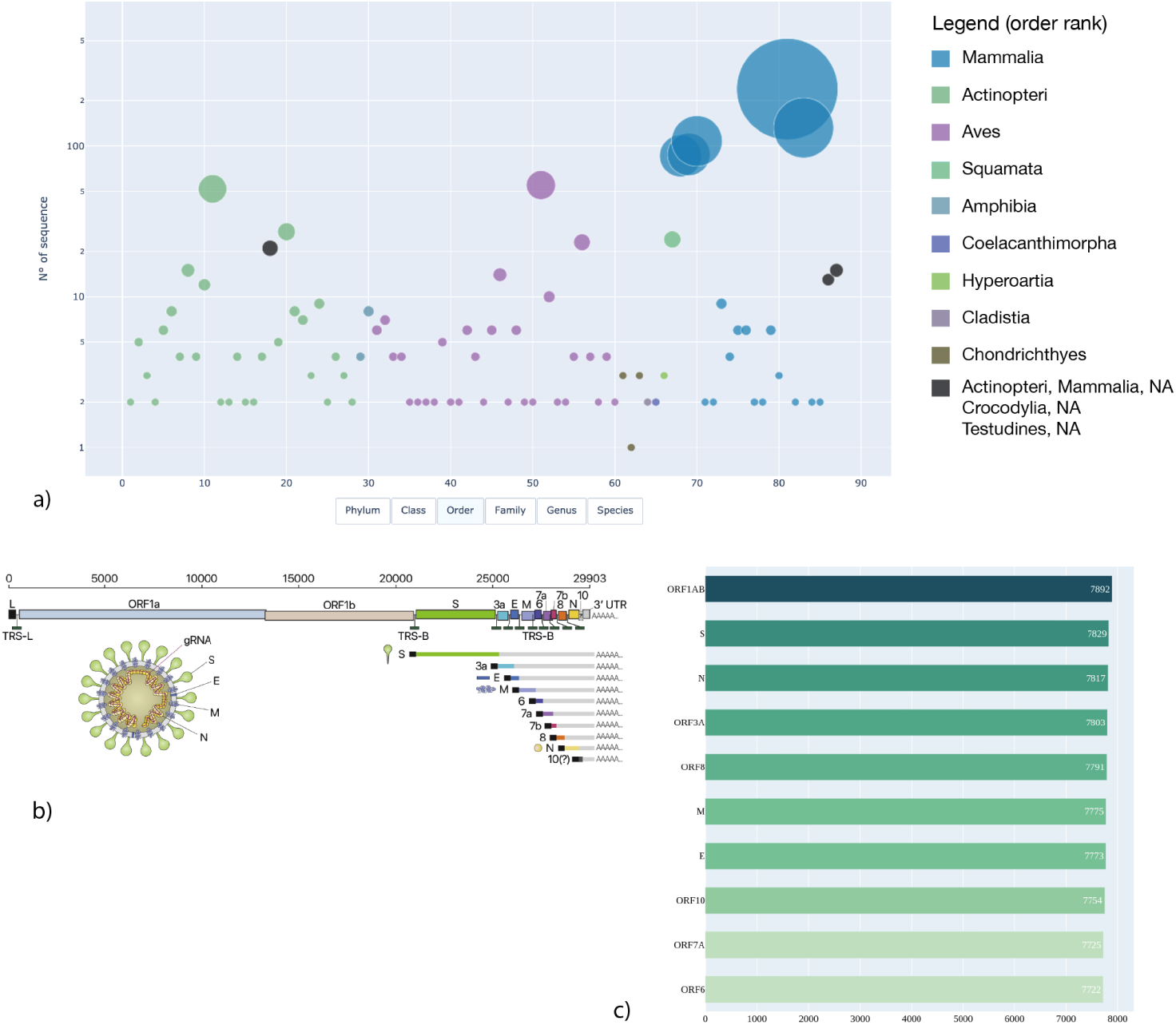
4a) Scatter plot of ACE2 accessions representing sequence abundances among taxa at order level; 4b) *SARS-CoV-2* representation, from [32]; 4c) gene distribution across accessions with complete ‘gene’ tag of *SARS-CoV-2* data.

Lastly, we explored the data available for *SARS-CoV-2* (Figure 4) using the following query “txid2697049” (where 2697049 is the specific severe acute respiratory syndrome coronavirus 2 TaxID; 29 of June, 2020). We obtained a total of 8,137 accessions. The top ten genes retrieved are shown in Figure 4c. In particular, the number of genes detected is quite similar among the top ten datasets and this is probably due to a high collection of genomes deposited into the database. The three most represented genes in the database are: ORF1AB (7892), followed by two important structural proteins: S (7829), the spike or surface glycoprotein, and N fragments (7817), the nucleocapsid protein. Considering the ORF1AB, several studies demonstrated its pivotal role among coronaviruses [55], providing a clinical target to break down *SARS-CoV-2* infection [31]. Regarding the second and third results, the nucleocapsid phosphoprotein is involved in packaging the RNA into virus particles and protects the viral genome. For these reasons, it has been suggested as an antiviral drug target [20, 60]. The spike glycoprotein, instead, is located outside the virus particle, mediating its attachment and promoting the entry into the host cell. It also gives viruses their crown-like appearance. In the very last research, the S protein was found as an important target for diagnostic antigen-based tests, antibody therapies and vaccine development [45, 53]. The entry of *SARS-CoV-2* is mediated by further processes, for example the activity of the protease TMPRSS2 [25]. Also in this case, the use of ExTaxsI can unearth similar proteases in possible intermediate hosts, revealing new insights into the mechanism of infection.

As also documented in Khailany et al., 2020 [31], the emergent and huge amounts of data collected in the last few months necessitates a large scale exploration of the data. The rapid increment of data releases may give some important insights about *SARS-CoV-2* behaviour in its host species, helping in improving not only our knowledge, but also models to predict COVID-19 outcomes and new drug targets.

## 3 Conclusions and future directions

ExTaxsI provides an easy-to-use standalone tool able to interact with NCBI databases and personal datasets, offering instruments to standardize taxonomy information and visualize vast quantities of data widespread on different taxonomic levels. It also provides interactive visualization plots, easily shareable through HTML formats.

The user-oriented interrogation of NCBI databases may help researchers involved in environmental genomics fields, from phylogeographic studies to DNA metabarcoding surveys, and also in projects related to the human health, as we demonstrated with the *SARS-CoV-2* case study.

With this work, we hope to meet the needs of a vast group of researchers, providing an instrument easy to install on common laptops and directly connected with NCBI databases. In our opinion, ExTaxsI data management ability with its visual interactive exploration can really improve the experimental design phase and the awareness of information available, facilitating the work and incentivizing data exploration and sharing.

## 4 Implementation

No specific system requirements are needed for the installation of ExTaxsI, however for the correct functioning of the software we suggest a minimum of 4GB of RAM. Moreover, to successfully run ExTaxsI, the following python libraries must be installed: NumPy, SciPy, Matplotlib, ipython, Pandas, SymPy, nose, genutils and Plotly, in addition to ETE toolkit [26]. To install all the dependencies compatible versions, we provide a requirement list at the GitHub page (https://github.com/qLSLab/extaxsi), with a detailed guideline to set directly a conda environment.

Regarding the organization of the tool, ExTaxsI is designed in separate modules, albeit interconnected, in order to work directly from different points of its workflow and to allow greater simplicity in the integration of additional modules in the future.

### 4.1 Database creation module

The module ‘Database’ allows users to create multi FASTA files composed of nucleotide sequences, taxonomic lists, genes names and their related accessions, starting from either a single query or a batch mode using csv/tsv files (Figure 1). After indicating the type of input, the tool asks, with the exception of the file accession, whether or not the user wants to integrate the query with one or more gene name/s (or other details). This step allows the user to restrict the research in NCBI if needed. In general, the output formats are i) a multi-FASTA file (widely used format for molecular sequences) and ii) text file in TSV format, with two columns composed by the accessions code followed by the taxonomy path of each accession at the six main levels separated by semicolons: phylum, class, order, family, genus and species. When requested by the user, the output file of genes names is in TSV format consisting of a table with two columns, one with the list of genes and the other with the frequency values of the respective genes found in the analyzed records. The tool also provides a summary table containing the most frequent genes from a list of taxid, accessions or organisms. In addition, it is possible to create a barplot with the top ten of this summary table, downloadable as a PNG file.

### 4.2 Visualization and statistical module

The module ‘Visualization’ allows users to create interactive plots, starting from Database module output or external sources (Additional files 3, 4 and 5) containing taxonomic lists. Before producing the plots, a dialogue box will ask the user to choose a filter value on the data based on the frequency. If the chosen filter value is 0, the tool processes all the data. Otherwise, all the taxonomic units that have not reached the minimum value are inserted into an additional text file, specifically created with a name containing the filter used. The available plots generated by ExTaxsI are i) ScatterPlot (Additional file 3), ii) SunBurst (Additional file 4) and iii) world map plot (Additional file 5). All figures created by the Visualization module can be downloaded as HTML format files. In detail, ScatterPlot uses taxonomy as input to produce a graph that indicates the quantity of each individual taxonomic unit; the interactive plot enables the user to: i) choose the taxonomic level to be displayed using the buttons located under the graph; ii) hover over points to show details, such as the number of records within taxa, names of selected taxa and name of the higher taxon from which they derives. The plot allows also to compare more data on mouse-over, highlight an area of interest with zoom function and view a specific group or remove taxa from the graph. SunBurst, instead, from a taxonomy input creates an Expansion Pie that allows to explore taxonomy by clicking on the taxonomic group of interest and showing the underlying taxa within a new SunBurst. Also in this case, hovering over points shows the number of records within taxa. Regarding world map plot, the initial input is processed in order to obtain geographic data. The tool exploits the ‘Country’ metadata stored in the NCBI records to produce a map indicating the position of each entry. In this step, based on the type of geographic data obtained, ExTaxsI divides results into two different arrays: i) a specific array of coordinates (if the coordinates are present in the record) or ii) a specific array of states names (if the coordinates are not present in the record). Also external sources can be processed and added to the map. In each map created, coordinates are indicated by green X signs, while States by red circles. Thinking of multiple taxa plotting, each symbol can have a legend that summarizes the data downloaded with the same country or coordinates description. Further, it is possible to see both genes and counts available among the accessions represented.

### 4.3 Taxonomy ID converter module

This module allows to convert TaxID to the main six ranks taxonomy and vice versa (phylum, class, order, family, genus and species); it can convert single manual inputs or multiple inputs from a tsv/csv file complete of a TaxIDs list.

## 5 Availability of source code and requirements

- Project name: ExTaxsI
- Project home page: https://github.com/qLSLab/extaxsi
- Operating system(s): Platform independent
- Programming language: Python
- License: GNU GPL version 3

### 5.1 Additional Files

**Additional file 1**: Gene list in TSV format obtained through ExTaxsI for the species *Gadus morhua*. Gene counts were extracted by 366,963 accessions (query: “txid8049[ORGN]”; 18 of June, 2020).

**Additional file 2**: World map plot in HTML format created via ExTaxsI extracting the values of ‘Country’ tag contained into 366,963 accessions of *Gadus morhua* (query: “txid8049[ORGN]”; 18 of June, 2020). Coordinates are indicated by green X signs, while States by red circles.

**Additional file 3**: ScatterPlot in HTML format created via ExTaxsI extracting the taxonomy of 388,603 accessions of *Gadiformes* Order (txid8043[ORGN]”; 22 of June, 2020).

**Additional file 4**: SunBurst Plot in HTML format created via ExTaxsI extracting the taxonomy of 388,603 accessions of *Gadiformes* Order (txid8043[ORGN]”; 22 of June, 2020).

**Additional file 5**: ScatterPlot in HTML format created via ExTaxsI extracting the taxonomy related to 1,189 accessions of ACE2 genes belonging to the *Vertebrata* taxonomic group (query: “txid7742[ORGN] AND ACE2[gene]”; 28 of June, 2020).

**Additional file 6**: SunBurst Plot in HTML format created via ExTaxsI extracting the taxonomy related to 1,189 accessions of ACE2 genes belonging to the *Vertebrata* taxonomic group (query: “txid7742[ORGN] AND ACE2[gene]”; 28 of June, 2020).

## 6 Declarations

### 6.2 Author’s Contributions

Giulia Agostinetto: Conceptualization, Investigation, Software development, Visualization, Original Draft Preparation, Review & Editing. Anna Sandionigi: Conceptualization, Original Draft Preparation, Review & Editing, Supervision, Project Administration. Adam Chahed: Software development, Visualization. Alberto Brusati: Investigation, Software development, Visualization, Review & Editing. Elena Parladori: Software development, Visualization. Bachir Balech: Review & Editing, Validation.

Antonia Bruno: Review & Editing, Validation. Dario Pescini: Review & Editing, Supervision. Maurizio Casiraghi: Funding Acquisition, Supervision. All authors read and approved the final manuscript, contributing critically important comments.

## 6.1 List of abbreviations

SILVA: High quality ribosomal RNA databases;
BOLD: Barcode of Life Data System;
UNITE: Database and sequence management environment centered on the eukaryotic nuclear ribosomal ITS region;
ETE: Environment for Tree Exploration;
QIIME2: Quantitative Insights Into Microbial Ecology;
FASTA: Text-based format for representing either nucleotide sequences or peptide sequences;
TAXID: Taxonomy ID;
HTML: Hyper-Text Markup Language;
COI: Cytochrome Oxidase I;
COI: Cytochrome Oxidase II;
COIII: Cytochrome Oxidase III;
CYTB: Cytochrome B;
ND2: NADH dehydrogenase 2;
ACE2: Angiotensin-Converting enzyme 2;
RBD: Receptor-Binding Domain;
PNG: Portable Network Graphics;
NCBI: National Center for Biotechnology Information;
ENA: European Nucleotide Archive

## 7 Acknowledgements

Many thanks are due to ELIXIR Biodiversity community for all the support.

